# Intra-slide calibration technology improves immunohistochemical harmonization within and between anatomic pathology laboratories

**DOI:** 10.64898/2026.06.04.730099

**Authors:** Glaucia Maria de Mendonça Fernandes, Wesley Wang, Anil Parwani, Saman S. Ahmadian, Michele Joana Alves, Joanna J. Philips, Jose Javier Otero

## Abstract

The reproducibility of immunohistochemistry in tumor tissue analysis across reference labs remains a persistent challenge. We tested the extent to which an intra-slide calibration technology mitigated discprepencies in inter-laboratory assays of p53 immunohistochemical (IHC) reactions in brain biopsies of glioblastoma (GB), IDH-wildtype. Intra-slide calibration technologies apply a 0-100% concentration scale incorporating primary surrogate and secondary antibodies to generate a standardized curve for DAB precipitation. IHC from GB samples was performed independently by pathology departments from two different hospital laboratories and were digitalized at 40x magnification using Aperio Image Scope software. Feature extraction, including intensity and texture parameters was performed using the *EBImage* package in R, followed by UMAP dimensionality reduction and DBSCAN clustering analysis. Our results show significant differences in intensity and texture clustering patterns between laboratory tissue samples and intra-slide calibration technology ruler caused by the different laboratories. Intra-slide calibration technology coupled with polynomial regression analysis improved ~90% the data harmonization. Our findings demonstrate a key role for computational pathology using intra-slide calibration technology to enable intra-laboratory consistency and inter-laboratory reproducibility. These advances strengthen the reproducibility of diagnostic assessments and support more objective, data-driven decision-making in neuro-oncology.

## Introduction

Immunohistochemistry (IHC) remains unparallelled in anatomic pathology due to its ease of deployment, technology penetrance in tertiary care and community hospitals, clear reimbursement rules that enable labs to make cost-based decisions on in-sourcing versus out-sourcing IHC-based assays, and for use as a rapid turnaround surrogate for a subset of molecular alterations ^1–7^. Primary brain tumor patient workflows include logistical challenges for patients such as coordination and transportation among radiologic imaging facilities, medical neuro-oncologists, radiation oncologists, and neurosurgical oncologists. Therefore, neuropathologists must proceed rapidly and accurately to test for molecular alterations to enable timely administration of radiotherapy and/or chemotherapy. Within neuropathology, the utilization of IHC-based surrogates for molecular testing has been widely adopted. Consequently, high-quality and reproducible IHC analyses across laboratories are vital for consistent diagnostic interpretation and clinical decision-making ^8–12^.

While the clinical importance of IHC is clear, reproducibility remains a major challenge. Variability in cold ischemia time, tissue fixation, antigen retrieval, antibody (AB) dilution, incubation times, detection chemistry, and chromogen development often leads to significant inter- and intra-laboratory discrepancies. These discrepancies may compromise not only diagnostic precision but also the reliability of prognostic and predictive IHC biomarkers ^4,13–22^. Furthermore, this heterogeneity remains a limitation to the development and validation of computational pathology and artificial intelligence (AI)-based tools for clinical decision-making ^5,23,24^. Although new approaches for standardizing and calibrating IHC assays have been proposed ^16–18,25–28^, the development of easily deployable technologies with minimal switching costs has eluded us. Quality assurance guidelines, such as those from the College of American Pathologists (CAP) and the Histochemical Society, emphasize that reproducible IHC results require internal and external controls to ensure comparability of staining intensity and interpretation ^29,30^. Intra-slide calibration technologies aim to fill this gap by acting as an IHC “ruler” to enable result harmonization between labs ^22^.

Intra-slide calibration involves printing arrays of co-resident control targets arranged explicitly as a 0–100% antigen-density scale (including primary- and secondary-target gradients). During IHC, the assay antibodies bind these on-slide targets to generate a calibrated 0–100% color-density (e.g., DAB) ruler, enabling reproducible, slide-specific quantification ^31,32^. This quantitative intra-slide reference can be incorporated into each staining run, enabling normalization of signal intensity across batches and laboratories. These intra-slide references can be applied to regular glass slides routinely utilized by anatomic pathology laboratories.

We aimed to evaluate the extent to which intra-slide calibration technology and computational pathology approaches could effectively reduce inter- and intra-laboratory variability in IHC staining of p53 in glioblastoma samples. Using a polynomial regression algorithm trained on the intra-slide calibration technology antibody ruler, we modeled the relationship between antibody concentration as a function of DAB staining intensity. These curves were then applied to clinical GB cases to assess their ability to align staining intensities across laboratories. We found that integration of intra-slide calibration technology with computational modeling can substantially improve IHC reproducibility and harmonization, thereby enhancing the reliability of p53 quantification in glioblastoma. Ultimately, such an approach could facilitate the development of standardized datasets for digital pathology, AI-assisted diagnostics, and automated image analysis, paving the way for more consistent assessment of clinically actionable biomarkers from IHC assays.

## Materials and Methods

### Samples collection and processing

Tissue samples from 15 patients with IDH-wildtype glioblastoma at the Ohio State University was identified, and two serial sections per case were prepared under an approved institutional protocol approved by Ethics Committee of The Ohio State University (IRB 2020C0150). Immunostaining was independently performed at Ohio State pathology department for sample processing (Lab 1), while the other slide was sent to the University of California San Francisco NIH Brain Tumor SPORE Biorepository for sample processing (Lab 2), using adjacent tissue sections to minimize biological variability. All slides were sectioned at 5 µm thickness using standard histological procedures. Although efforts were made to maintain comparable pre-analytical conditions, including routine clinical handling and processing workflows at both institutions, variations in slide preparation, storage duration between sectioning and staining, and local laboratory conditions may have contributed to inter-laboratory variability and reflect real-world practice. Immunohistochemistry for p53 antibody was performed in both laboratories in slides with Dako clone DO7, 1/1000 dilution, following their respectively validated clinical protocols for antigen retrieval, antibody dilution, incubation times, detection chemistry, and chromogen. TP53 mutational status by sequencing was not provided by the IRB. Whole slide images were obtained and digitalized at 40x magnification using Philips Ultra-Fast Scanners (Koninklijke Philips N.V.). For downstream analysis, 10 tiles per sample were extracted for each case, while for the intra-slide calibration technology (I-SCT), each concentration level was captured in its entirety to ensure full signal representation. Image analysis was conducted using Aperio ImageScope software (v12.4.3.5008, Leica Biosystems).

### Intra-slide calibration technology

The intra-slide calibration technology used in this study is based on the Process Record Slide (PRS) IHC controls (PRS; https://ihc-prs.com), which function as certified on-slide process calibrators and provide a rigorously standardized alternative to conventional tissue-based controls by incorporating reference targets with defined biochemical properties ^31,32^. Each PRS slide features printed primary and secondary protein targets arranged in predefined antigen-expression gradients spanning a 0–100% scale, including both positive and negative controls. During IHC staining, the assay’s primary and secondary antibodies bind to these co-resident targets, and subsequent chromogenic development with DAB produces a calibrated 0–100% color-density curve that faithfully captures the actual staining performance of that individual slide. This calibration ruler serves as an absolute quantitative reference for stain-intensity assessment and provides the foundation for our downstream inter-slide harmonization pipeline, which corrects staining variability across slides, batches, scanners, and laboratories. **Fig. 1** illustrates the inclusion of the Intra-slide calibration technology and our computational workflow highlighting how this ruler could contribute to standardized and classifying the cells by intensity measure.

**Figure 1.**
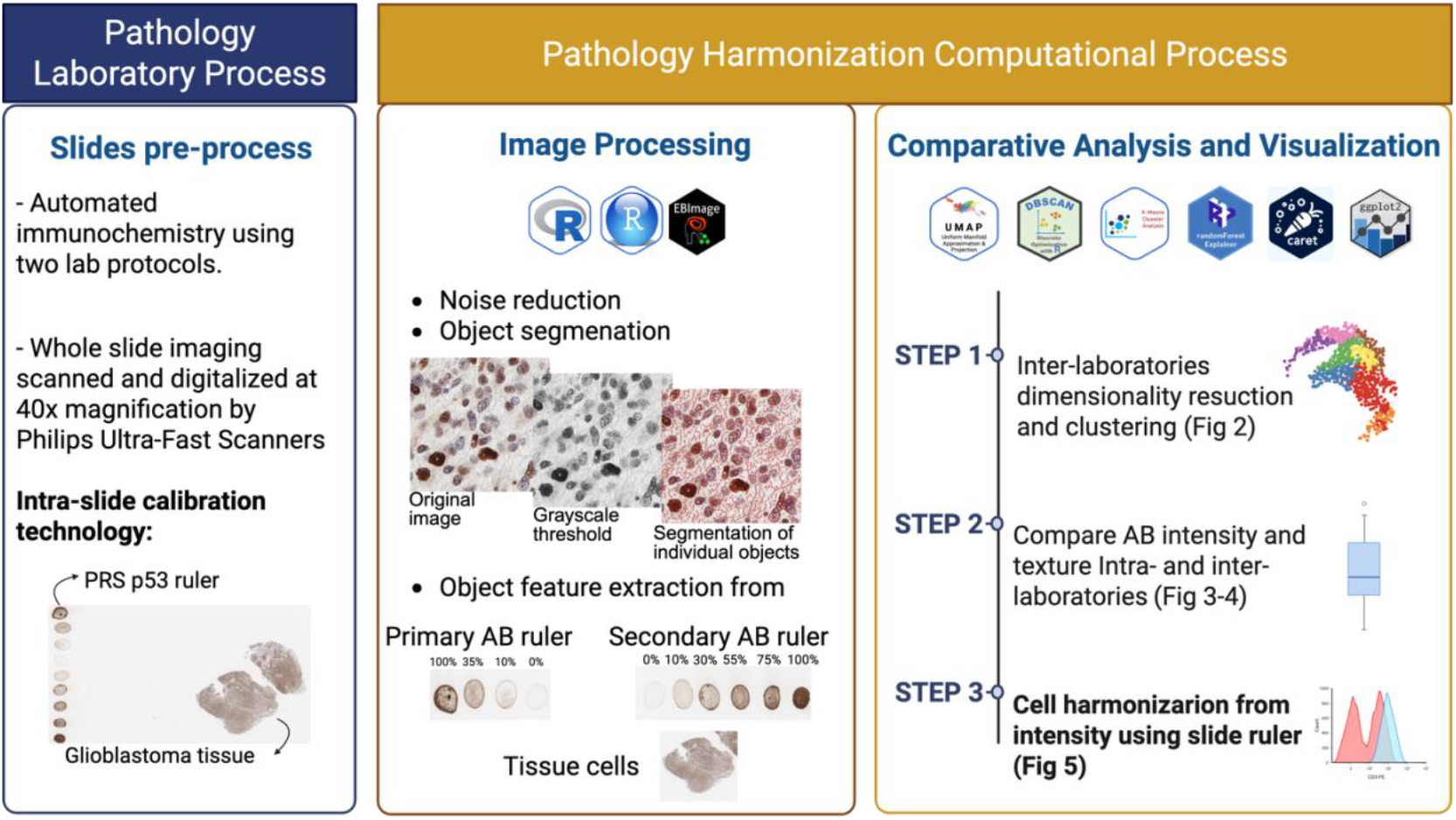
Computational workflow for pathological slide analysis integrating immunohistochemistry (IHC) data and R-based image processing using antibody intensity scales with Intra-slide calibration technology. The pipeline begins with slide preparation following automated laboratory immunostaining protocols, followed by digital scanning. R packages such as EBImage, PCA, DBSCAN, and UMAP are used to perform image processing and feature extraction on slides stained with both primary and secondary antibodies. Mean intensity values are used to construct antibody-specific intensity scales. Dimensionality reduction (via PCA and UMAP) and clustering (via DBSCAN) support comparative analyses and classification tasks. The workflow addresses three main steps: (1) comparing cell intensity, texture, and shape heterogeneity across laboratories for p53 IHC; (2) evaluating consistency between primary and secondary antibody scales across cases and laboratories; and (3) harmonizing cell intensity based on slide concentration ruler. Data visualization is performed using ggplot2 and plotly, with outputs presented as scatter plots, UMAP projections, and bar plots to illustrate inter-laboratory differences.

### Pathology Computational Workflows

#### Image analysis

For each case, the same representative regions of interest (ROIs, 240 x 240 pixels) were manually selected from WSI processed in both laboratories. For the intra-slide calibration technology, each concentration level was captured the across the entire field to ensure complete signal representation. All images were exported as TIFF images for quantitative image analysis.

Image pre-processing and feature extraction was performed in R software using *EBImage* package ^33^. Each tile was first converted to an RGB array and normalized to correct illumination and intensity bias. Background noise was reduced using a Gaussian filter (σ = 1-4), followed by contrast enhancement to standardize signal dynamic range across images. To isolate the DAB (3.3’-diaminobenzine) chromogenic signal, color images were converted into grayscale intensity maps using the *channel* function (1-gray), highlighting the higher pixel intensity values of the DAB-positive regions. Segmentation of DAB-positive areas was achieved using adaptative local thresholding (*tresh* function for window size = 25×25 pixels, offset = 0.05), improving segmentation accuracy in heterogeneous staining conditions ^34^. We implemented minimal morphological refinements by applying hole-filling operations (*fillHull* function) to internal gaps within stained nuclei, followed by connected-component labeling (*lwlabel* function) to assign unique identifiers to each segmented object, enabling robust delineation of individual immunopositively cells even in regions with heterogeneous staining or partial background artifacts common in IHC tissue sections.

For each segmented object, a comprehensive set of morphometric and intensity-based features were computed using EBImage function *computeFeatures*.*basic, computeFeatures*.*moment, computeFeatures*.*haralick*. Extracted descriptors included object area, perimeter, circularity, centroid coordinates, mean and integrated optical density, and texture descriptors ^35^. Feature matrices were concatenated across red, green and blue color channels (RGB) to create per-tile datasets, which were aggregated to generate case-level feature summaries such as positive area fraction, mean intensity and heterogeneity indices. All processing steps were implemented through reproducible R scripts to minimize operator bias and ensure batch consistency. The tile-based approach allowed detailed local quantification of staining patterns while maintaining computational feasibility for large sample sets.

#### Inter-lab tissue heterogeneity

To compare the staining patterns and cell populations across laboratories, extracted features from the cell nuclei underwent dimensionality reduction using Uniform Manifold Approximation and Projection (UMAP) implemented through the *umap and uwot* package ^36^. This nonlinear dimensionality reduction technique enables visualization of multidimensional features distribution in a two-dimensional embedding. We also performed density-based spatial clustering of application with noise (DBSCAN) using *dbscan* package, to identify distinct cell clusters based on feature similarity ^37,38^. Each cluster was characterized by staining intensity, textural, and morphological features. We then used the *c*_*x*_ and *c*_*y*_ coordinates to identify the cell closest to the cluster centroid, which was selected as a representative example of that cluster.

#### Intra- and inter-lab antibody concentrations rule heterogeneity

To compare the staining patterns in the concentrations of the Intra-slide calibration technology across laboratories, the intensity and texture features extracted were analyzed using Principal Component Analysis (PCA) to reduce dimensionality and remove collinearity among correlated variables ^39^, followed by Random Forest algorithm to test the accuracy in the reproducibility. Dispersion and box plots were generated using the *ggplot2* and *ggpubr* packages. These visualizations were used to compare the distributions of intensity-texture-related features between laboratories.

#### Cell tissue intensity harmonization

To harmonize staining intensity across samples and laboratories, a cell-level harmonization procedure was implemented using the antibody ruler generated from the intra-slide calibration technology slide. First, all features extracted from the Intra-slide calibration technology concentrations and from the nuclei objects underwent PCA. The intra-slide calibration technology served as an antibody ruler, where a standardized intensity curve based on 0-100% antibody concentrations for the secondary AB was used as a reference for the calibration function. A polynomial regression model was then trained using the intra-slide calibration technology curve to fit the relationship between DAB optical density and antibody concentration ^40,41^. This model was applied to the cellular features obtained from tissue samples, allowing prediction and normalization of staining intensities according to the standardized scale from the intra-slide calibration technology.

#### Scale-invariant inter-laboratory comparison

Because pre-calibration (RAW) and post-calibration (PRED) intensity values are expressed in different units (principal component values approximately 0–1 vs calibrated percentages 0– 100%), we quantified inter-laboratory discrepancy using the standardized mean difference (SMD), which is invariant to the scale previously described ^42^. For each case, we computed and summarized ∣SMD∣ for RAW and PRED conditions.

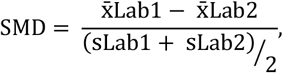

Where 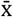 denotes sample mean and s denotes sample standard deviation. Lower ∣SMD∣ indicates greater cross-lab harmonization. We then derived per case Δ∣ SMD ∣ = ∣ SMD ∣⁄RAW − ∣ SMD ∣⁄ and a percentage reduction 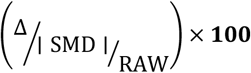. To avoid unstable ratios when the baseline discrepancy is trivial, we omitted the per case percent reduction whenever ∣ SMD ∣⁄RAW< 0.10 (prespecified baseline threshold). The global harmonization effect was summarized by the median ∣SMD∣ across cases before vs after calibration and reported as a percent reduction in the median. A priori sensitivity analyses confirmed that the global median based reduction is insensitive to the baseline threshold; only the per case percent column is suppressed when baseline ∣SMD∣ is near zero.

### Statistical analyses and software environment

All statistical and computational analyses were conducted in *R* software (v4.3.2) within the *RStudio* interface (v2023.09). Image processing, segmentation, and feature extraction were performed using the *EBImage* package (v4.38.0). Dimensionality reduction was performed using the *umap* package (v0.2.10.0), clustering analysis of cellular features using *dbscan* package (v1.1-12) and principal component analysis using *factoextra* package. Machine learning procedures, including random forest and polynomial regression modeling for AB concentration harmonization were conducted using *randomForest* package, *caret* package and base *stats* functions.

All quantitative variables were tested for normality using *Shapiro*.*test* and inter-laboratory comparisons were performed using two-tailed Student’s t-test. Statistical significance was set as p>0.05. Model performance was evaluated through accuracy, 95%CI, no information rate, kappa and p-value metrics derived from confusion matrix.

Data visualization was conducted using *ggplot2* (v3.5.1), *ggpur* (v0.6.0), and *patchwork* (v1.2.0) to generate violin, box, bar and density plots for comparative feature distributions before and after harmonizations. All imaging data only upon request to the correspondent author.

## Results

### Inter-laboratory heterogeneity in tissue cell immunohistochemistry preparation and analysis

To test the efficacy of intra-slide calibrate technology, we chose p53 immunohistochemistry of brain tumors. The p53 immunohistochemical reaction appears nuclearly in cells, and hereon we refer to these as p53 signals. To achieve this, we use intra-slide calibration technology in independently automated IHC standing from Lab 1 and Lab 2. Cell signal was visibly different between labs as seen in the two laboratories as an example of case 1 on **Figure 2A**. After the whole slide imaging, digitization was performed in the same scanner system followed by extraction of image intensity, image texture and nuclear morphology features **(Fig. 2A)**. The UMAP plots from intensity and texture features revealed that most cells in the tissue show significant differences in imaging features as they are clearly separated across labs for each specific case, with minimal overlap between Lab 1 and Lab 2 cell populations (note the lack of overlap between Lab 1 and Lab 2 nuclei in **Fig. 2B and SFig. 1)**. This results highlights the impossibility to standardize, even within same lab, given DAB chemistry, batch issues.

**Figure 2.**
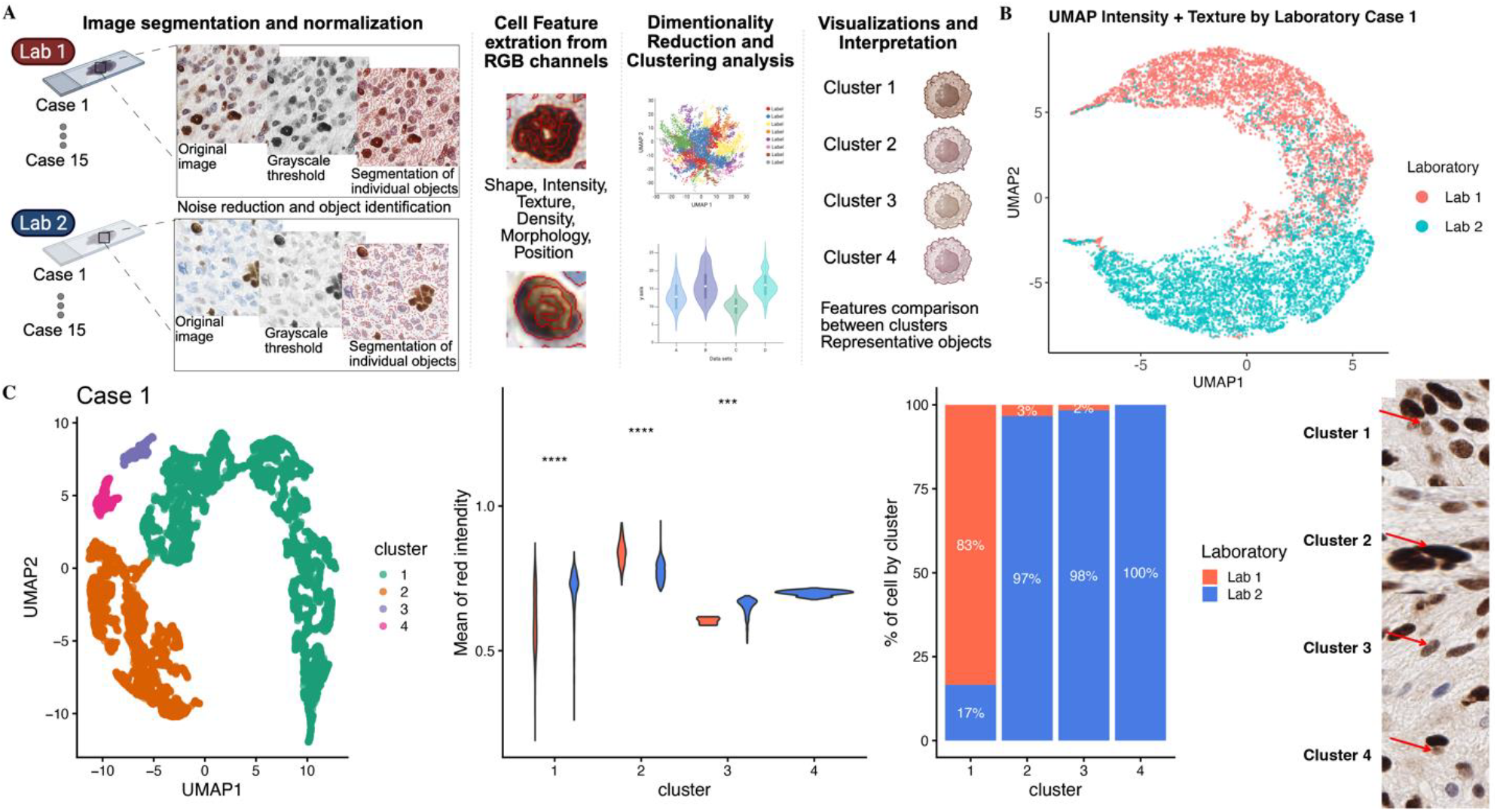
Analysis of Inter-Laboratory Variability in Image Processing and Feature Extraction. (A) workflow for the 10-tile selection from two different laboratories (Lab 1 and Lab 2), followed by image processing (grayscale conversion and segmentation), and feature computation (intensity, texture, shape, and moment) using EBImage R package, and dimensionality reduction using UMAP for visualization. (B) UMAP plot presents the distribution of intensity and texture features for 15 different cases between the laboratories. Samples from Lab 1 are shown in red, and samples from Lab 2 are in blue. (C) Each set of plots represents the distribution of cell morphology by DBSCAN clusters for specific cases, visualized in UMAP plots showing the cluster distributions (left), violin plots displaying the clusters intensity differences between laboratories (middle left), bar plots in left showing the % of cells of each lab in each cluster (middle right), and representative cell images for each cluster (right). Samples from Lab 1 are in red, and samples from Lab 2 are in blue. Violin plot representing the differences between laboratories, performed by T test and present statistical significance: ns: p > 0.05; *: p <= 0.05; **: p <= 0.01; ****: p <= 0.0001.

To specifically assess the cell intensity-related effects, we performed a second UMAP embedding based on cell intensity followed by clustering by the DBScan method. These identified clusters are based principally on cell intensity features. We visualize the differences of cell intensity and the proportion of detected cells in each cluster with violin and box plot+ in **Figure 2**. Representative examples of cells from each cluster visually corroborated these quantitative findings, evidencing consistent variations between laboratories in the cell intensity distribution for each case (**Fig. 2C and SFig. 2)**. Also, we found similar clustering outcomes by all extracted imaging features including intensity, texture and shape **(SFig. 3)**. In summary, significant inter-laboratory variability in the distribution patterns of the cell imaging features between the laboratories was noted. Although these two tests were validated using appropriate CAP-compliant standard operating procedures, significant variation nevertheless persisted. This variation may be attributed to differences in sample preparation and processing techniques inherent to the limitations of CAP-compliant methodologies, highlighting the importance of on slide or off slide control run by same lab and computational pathology workflows to enable standardizations that ensure reproducibility and reliability of diagnostic results.

**Figure 3.**
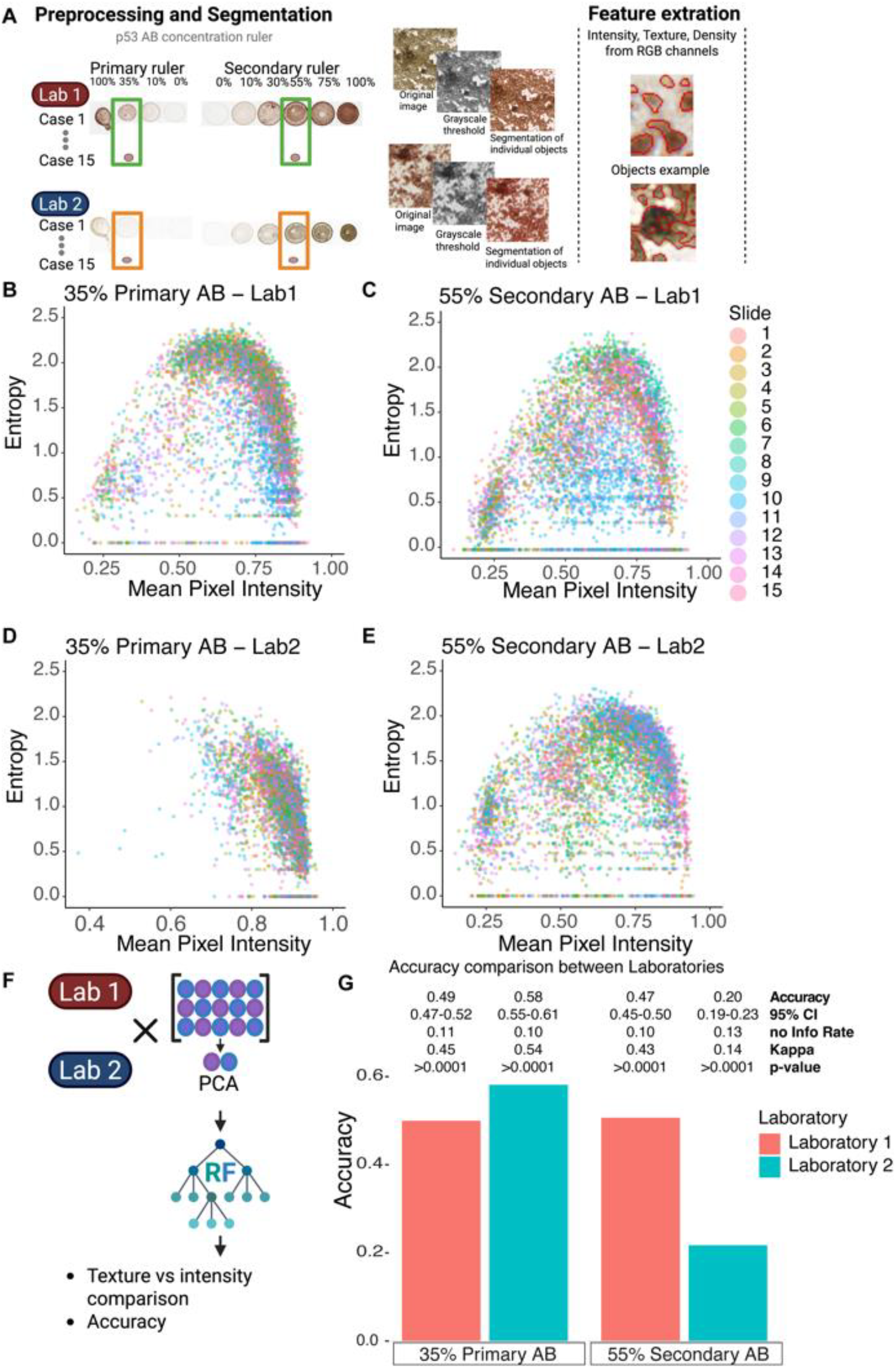
Intensity and texture analysis Intra-laboratory. (A) Workflow showing preprocessing of image extract features from selected objects in the scale. Graphs for mean Intensity and entropy (texture) distribution across all cases for the 35% Primary antibody concentration from (B) Lab 1 and (D) Lab 2, and for the 55% Secondary antibody concentration from (C) Lab 1 and (E) Lab 2. Objects are color-coded by pathology case. (F-G) Workflow and Bar plot presenting the differences between the model accuracy for each antibody concentration.

### Intra-lab heterogeneity in Intra-slide calibration technology immunohistochemistry preparation

We next sought to evaluate the capacity of each laboratory to reproduce the p53 immunostaining result from primary and secondary antibody concentrations on each pathology case. To achieve this, we performed homogeneity testing using image segmentation and image feature extraction of objects derived from the slide scale ruler at the 35% and 55% secondary AB concentration. **Figure 3A** illustrates the image segmentation approach and samples of segmented objects from the slide scale ruler. Using scatter plots of the texture feature Entropy as a function of Mean Pixel Intensity of each segmented object at 35% and 55% secondary Ab concentration we observed distinct distribution patterns between laboratories, with variable dispersion and clustering of objects across cases (**Figure 3 B-E)**. Notably, **Figure 3D** demonstrates increased homogeneity compared to **Figure 3B**, with tighter clustering and a more consistent relationship between intensity and entropy. In contrast, **Figure 3B** exhibits broader dispersion and less defined structure, indicating greater variability in staining response. Similar trends are observed for the secondary antibody conditions **(Figure 3C and 3E)**, where differences in clustering patterns further highlight inter-laboratory variability. Together, these results indicate that both intensity and texture features vary substantially intra- and inter-laboratories, impacting the reproducibility and comparability of IHC measurements.

These data were used to model image features as a function of secondary AB concentration with a random forest algorithm. Next, we were interested in determining the extent to which a random forest classifier could determine from which case each segmented object came from. To this end, we separated 70% of the segmented objects for training and 30% as test objects. In this approach, a lower accuracy of determining the slide from which the object indicates increased homogeneity across samples (see workflow in **Figure 3F**). Lab 2 had greater intra-laboratory reproducibility (note accuracy of Lab 2 versus Lab 1 in **Figure 3G**). We conclude that intra-slide calibration technology can be utilized to assess the reproducibility of IHC assays within one laboratory. We further conclude that intra-slide calibration technology can be used to compare the reproducibility of different laboratories.

### Inter-lab heterogeneity in Intra-slide calibration technology immunohistochemistry preparation

By mere visual inspection, the results of p53 IHC reactions on the slide calibration are quite distinct despite similar staining protocols. Using a similar homogeneity testing approach as described above, we next determined the extent of inter-laboratory variability in p53 immunostaining by evaluating the imaging features of the slide ruler. The workflow is illustrated in **Figure 4A**. To evaluate the differences between all objects of each scale, we projected each object onto 2-Dimensional space using the UMAP algorithm (**Figure 4B**). We note that each concentration gradient shows well-formed clusters with the objects of the same concentration but different labs occupying very different areas on the UMAP. This data implies that significant variance is noted between the labs on the scales. To achieve the difference between specific concentration in the scale, we compared the 35% from primary and 55% from secondary AB concentrations from the intra-slide calibration technology across the laboratories. Scatter plots of segmented objects are shown in **Figure 4C** with each object color coded by laboratory, and which shows significant difference between labs. To further test this, we performed homogeneity testing with a random forest classifier. We separated 70% of the segmented objects for training, and 30% as test objects. The accuracy of the trained random forest classifier on the test set was then recorded and plotted in. In our random forest classifier, we predict from which lab the object derives from as a function of image features of the extracted objects. We note very high accuracy in detecting from which laboratory the objects derived from 35% primary antibody (Acc=82%, p<0.001) and 55% secondary Antibody (Acc=79%, p<0.001), indicating low homogeneity, and therefore, high variability between labs (**Figure 4D)**.

**Figure 4.**
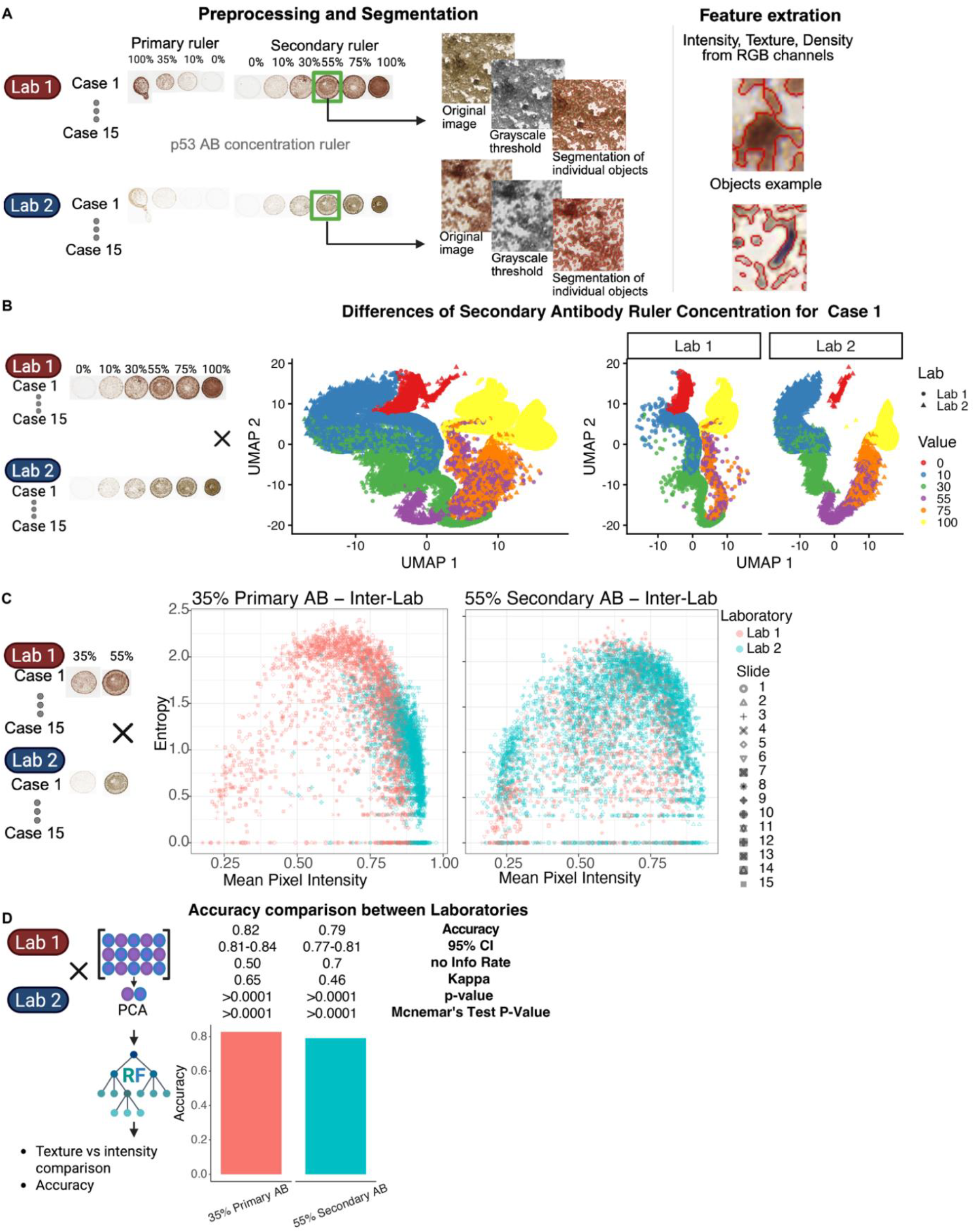
Intensity and texture analysis inter-laboratory. (A) Workflow showing example preprocessing images to extract the feature from all objects in the scale. (B) UMAP plot of the case 1 clustering the differences between each concentration value of the Secondary antibody in the intra-slide calibration technology across laboratories (item 2). (C) Graphs for mean Intensity and entropy (texture) distribution across all cases comparing the Lab 1 with Lab 2 for the 35% Primary antibody concentration, and for the 55% Secondary antibody concentration (item 1). (D) Workflow and Bar plot presenting the differences between the model accuracy for each antibody concentration.

### Computational harmonization across laboratories of p53 Antibody Intensity using Intra-slide calibration technology

As p53 testing can be used as a proxy for molecular testing, it is critical that across laboratories a harmonization of the IHC data between labs is necessary. To achieve this, we created a computational workflow **(Fig 5A)** based on the secondary AB 0 to 100% scale as a ruler in the slides. Mean pixel intensity data from red, green, and blue color channels were extracted from the segmented images of the slide ruler and the p53 immunolabelled cells and underwent PCA to obtain the value for PC1. We then generated a polynomial regression where secondary AB concentration was modeled as a function of mean pixel intensity PC1 value. This model was then used to predict the PC1 value for each slide in the different laboratories. This workflow harmonizes the laboratories results of the p53 cell intensity quantification. We can observe a similar cell intensity between the laboratories after modeling. **Figure 5B** presents the density plot for the Case 1 harmonization across labs before and after our workflow prediction (Cases 2 to 15 in **SFig. 5). Figure 5C** presents a comparison between labs showing all cases, the first bar plot presents the PC1 data of mean intensity while the second bar plot presents the data from the polynomial prediction using the inter-slide calibrate technology. To compare percent reduction between the harmonization without unit artifacts, we summarized inter-lab discrepancy per case using the standardized mean difference for before (RAW: PC1, ~0–1) and after (PRED: calibrated %, 0–100) calibration, then we calculate the differences between the ∣SMD∣ calibration. Across cases, the median ∣SMD∣ for RAW was 1.66 compared to 0.16 for the PRED, a with a Δ∣ SMD ∣ of 1.5 or 90.29% reduction, indicating markedly improved cross-Lab harmonization **(Figure 5D; Supplementary Table S1)**. Our results demonstrate the innovative and promising approach using the intra-slide calibration technology to reduce the variation in the p53 antibody intensity quantification across the labs. These results enhance inter-laboratory reliability, reproducibility, improving interpretation for pathologists and enabling future automation in digital pathology diagnostics.

**Figure 5.**
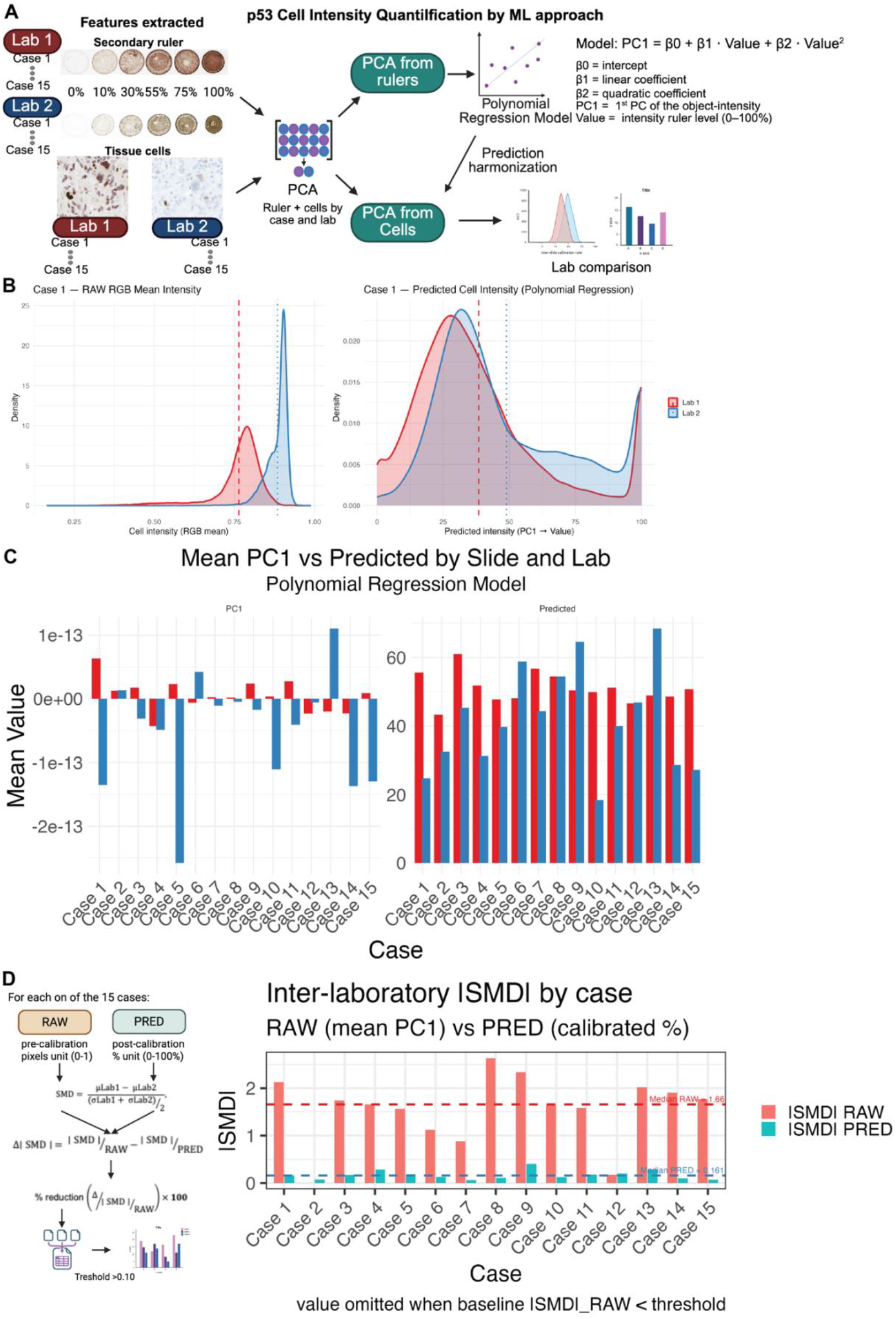
Predictive model to harmonize p53 antibody cell intensity within and between laboratories. (A) Schematic workflow of the harmonization process illustrating the extraction of intensity features from the tissue samples and the antibody concentration ruler, followed PCA analysis to capture the major sources of variation in intensity and reduce dimensionality before harmonization. The polynomial regression algorithm was applied to the secondary antibody ruler and used to harmonize the tissue-derived intensity cells values. (B) Density plot displaying the distribution of raw cell intensity vs predicted cell intensity across laboratories, based on the secondary antibody ruler applied to each slide. (C) bar plots highlighting the differences of the raw intensity data for the PC1 (left) and the results after harmonization using the polynomial regression model (right). (D) Inter-laboratory SMD by case before (RAW; principal-component units) and after (PRED; polynomial regression calibrated %) I-SCT. Lower values nearby to 0, indicate better cross-lab harmonization. Dashed lines mark the median SMD across cases in each condition. Percent reduction is reported per case only when the baseline SMD > 0.10; values below this threshold are omitted (see Methods). Abbreviations: SMD: standardized mean difference.

## Discussion

### Identification of antibody dilution in clinical immunohistochemistry

It is well recognized that the use of p53 IHC as a surrogate for TP53 mutation is challenged by inter-laboratory variability, which persists even under CAP-compliant validation. Differences stem from protocol and platform factors (antibody lot/dilution, retrieval and detection chemistry, interpretive variability) despite adherence to analytic-validation requirements. The 2024 CAP guideline update harmonizes validation across predictive markers but does not specify a per-slide intensity calibrator for the IHC readout ^29^. This leaves the analytic readout without a slide-level quantitative anchor in routine practice.

The CAP framework is an important step toward standardizing laboratory practice, mandating pre-clinical validation/verification, setting a ≥90% concordance benchmark, distinguishing predictive from non-predictive assays, and defining triggers for revalidation. Nevertheless, important limitations remain. Several recommendations rely on expert consensus rather than high-quality empirical data, and the 90% concordance threshold may still allow clinically meaningful discrepancies, particularly for assays used to guide therapy. In parallel, external quality-assessment (EQA) programs (e.g., NordiQC, UK NEQAS ICC/ISH) repeatedly document cross-laboratory variability and identify suboptimal protocol choices, underscoring the need for methods that stabilize the analytic intensity itself ^13,29,43,44^. Both programs consistently report inter-site dispersion and protocol-sensitive performance, reinforcing the need for an analytic anchor. For predictive IHC, multicenter PD-L1 Blueprint studies show clone- and platform-dependent differences in tumor-cell positivity that shift clinical classification, indicating that appearance-level harmonization does not ensure analytic equivalence at cut-offs ^45,46^. These differences persist even when slides “look” similar, highlighting the need for quantitative readout control. Our findings are concordant, demonstrating measurable variability in cell-level features across laboratories, consistent with slide-to-slide differences in antibody–epitope response that persist even under CAP-compliant conditions. Such technical variation directly impacts interpretability and highlights an ongoing barrier to achieving truly consistent IHC results. Addressing these residual challenges is essential for ensuring diagnostic accuracy and optimizing clinical decision-making in neuro-oncology. The clinical relevance is particularly evident for p53, where recent reports show imperfect concordance between IHC patterns and TP53 status with both technical and interpretive contributors. This aligns with recent p53 studies that quantify discordance and attribute part of it to analytic variability ^44,47,48^.

### Intra-slide calibration for quantitative cell-level intensity and cross-laboratory harmonization in IHC

Current validation and EQA frameworks lack a quantitative, per-slide calibrator for IHC intensity (i.e., a practical “on-slide reference”) ^29,43,44^. This gap is not fully addressed by the current 2024 CAP guidelines ^29^. Additionally, persistent inter-laboratory variability documented by EQA programs (e.g., NordiQC; UK NEQAS ICC/ISH) reinforces the need for a slide-level analytic anchor ^13,43,44^. We address this gap by using the PRS as an on-slide 0–100% quantitative reference that anchors the IHC readout and applying our intra-slide calibration workflow that fits a global PCA step followed by slide-specific polynomial regression to map pixel values to a calibrated, assay-agnostic intensity scale. The resulting parameters are interpretable, transparent, auditable, and version-controlled. This workflow converts raw optical signal into a calibrated estimate of antibody–epitope interaction strength, enabling cross-slide and cross-lab comparability. This procedure mitigates analytic drift arising from retrieval, detection, and optical factors but cannot restore pre-analytic antigen loss. Applying the workflow across two laboratories with distinct processes, we observed a ~90% reduction in median inter-laboratory differences |SMD| and more stable classifications near decision thresholds particularly for markers with narrow analytic boundaries. Prospectively, calibration re-checks will be triggered by antibody-lot, retrieval/detection, platform, or environmental changes, with verification of fit quality, preserved monotonicity, and internal control tolerances, aligned with CAP revalidation principles ^29^. By reducing domain shift, intensity calibration reduces inter-laboratory variance and enhances inter-laboratory reproducibility and strengthens external validity of computational and AI models ^49^. Because the polynomial regression operates strictly within the observable intensity domain ^50^, it cannot reconstruct signal lost to ischemia or over-decalcification. In settings characterized by multifactorial variability and complex spatial artifacts. A complementary deep-learning residual strategy may further improve accuracy while preserving auditability and interpretability ^51^.

In neuro-oncology, small intensity shifts around practical boundaries for p53 can alter classification and downstream management; recent studies show imperfect concordance between p53 IHC patterns and TP53 alterations, with both technical and interpretive contributors ^47,52^. By anchoring intensity to a calibrated scale, the intra-slide ruler reduces boundary instability, supporting more consistent workflows and providing higher-quality inputs for AI models trained/validated on multi-site data. Beyond p53, the approach is applicable to biomarkers with low-expression cut-offs (e.g., PD-L1, HER2 0 vs 1+), where small analytic shifts have therapeutic implications. This is consistent with PD-L1 Blueprint comparability data and recent HER2 guidance on low expression ^43–45^. Future research should validate applicability across additional staining patterns and tissue types.

Our study involved a modest number of cases and laboratories, which may not fully represent the diversity of staining protocols encountered in broad clinical settings. While our results demonstrate improved harmonization within and between laboratories, larger multi-institutional studies will be essential to confirm the clinical impact of I-SCT-based harmonization on diagnostic reproducibility. Key endpoints include reclassification rates, inter-rater reliability (κ/ICC) near decision thresholds, and performance on EQA/proficiency-testing tissues ^13,29,43,44^. Although not applied here (single lot), calibration checks will be embedded in CAP-aligned change-control (new lot; retrieval buffer/pH; detector/platform updates) with pre-release verification of fit, monotonicity, and internal-control tolerances.

### Good Practice I-SCT implementation and recommendation

When feasible, we propose that laboratories should implement and document a per-slide analytic calibration of IHC intensity using I-SCT based methods on antibody-concentration ruler and a slide-specific polynomial response model; calibration should be included in the validation dossier and applied to clinical runs, complementing (not replacing) pre-analytic controls, EQA participation, and CAP revalidation triggers.

In conclusion, our computational intra-slide calibration-based harmonization workflow for IHC protocols provides a robust and scalable strategy to minimize technical variability and ensure reproducible staining within and between laboratories. By generating standardized and quantifiable intensity profiles, this approach enhances the reliability of diagnostic interpretation by pathologists and delivers consistent, high-quality data for the training and validation of artificial intelligence models in digital pathology. These advances strengthen the reproducibility of diagnostic assessments and support more objective, data-driven decision-making.

## Acknowledgments

The authors gratefully acknowledge the support provided by The Ohio State University for their institutional and research infrastructure support. We also thank Florida International University (FIU) for their continued support throughout this study. We acknowledge the support and collaboration Digital Pathology Team and Division of Molecular Pathology team from Department of Pathology at The Ohio State University Wexner Medical Center, and Brain Tumor SPORE Biorepository at University of California, San Francisco whose facilities, expertise, and resources were invaluable to the successful completion of this study. Workflow figures in this manuscript were created using BioRender.com platform.

## Author contributions statement

All authors contributed to the study conception and design. Material preparation and data collection was performed by GMMF, WW, JJO and JP. Data analysis and first draft of the manuscript was written by GMMF and JJO. All authors discussed and commented on previous versions and approved the final manuscript.

## Statements and declarations

### Ethical considerations

This study was performed in line with the principles of the Declaration of Helsinki. Approval was granted by the Ethics Committee of The Ohio State University (IRB 2020C0150).

### Consent to participate

Informed consent was obtained from all individual participants included in the study.

## Data availability statement

Imaging data only upon request to the correspondent author.

