## Supplementary Figures and Tables for "Intra-slide calibration technology improves immunohistochemical harmonization within and between anatomic pathology laboratories"

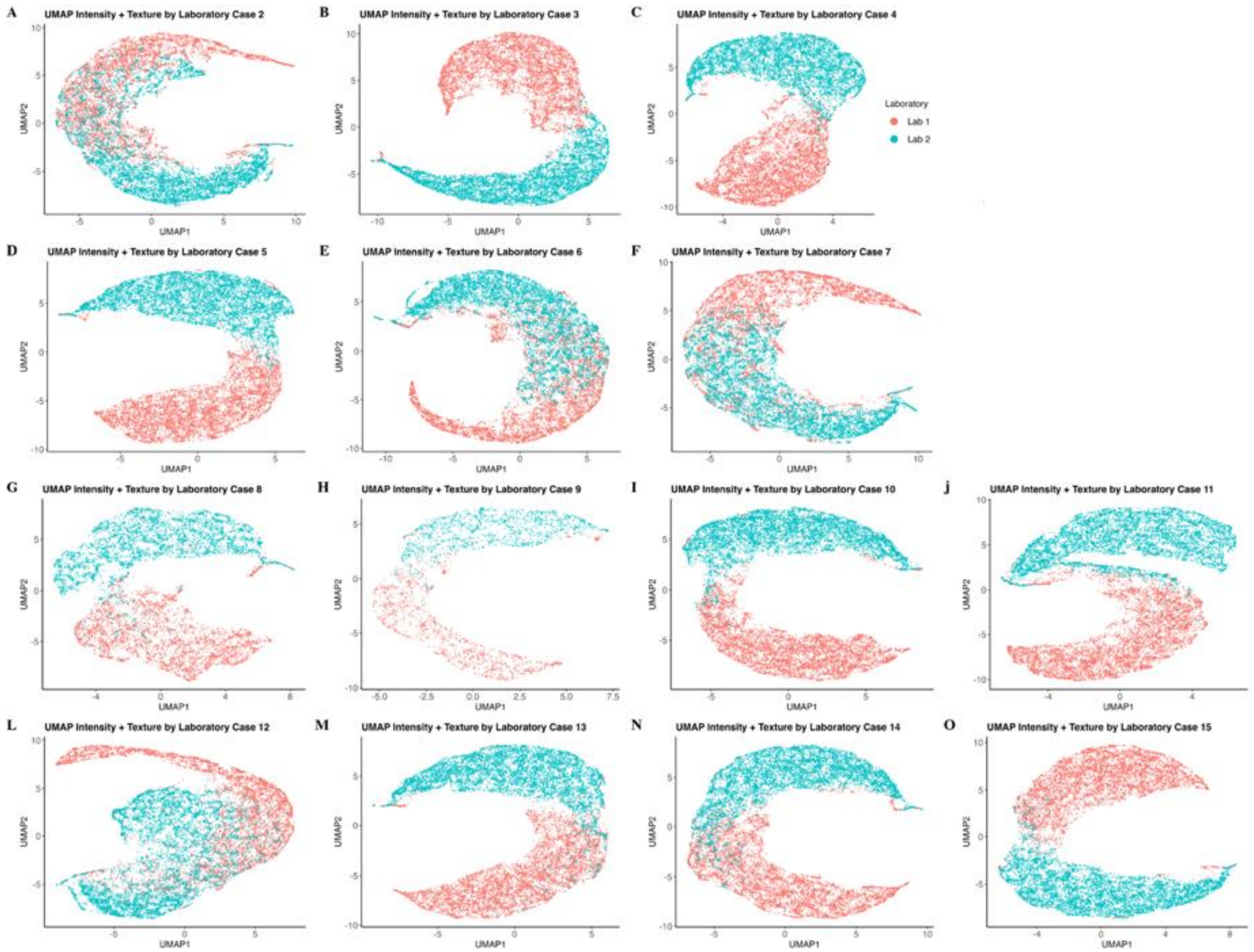


**Supplementary Figure 1: Analysis of Inter-Laboratory Variability in Image Processing and Feature Extraction.** (A-O) UMAP plot presents the distribution of intensity and texture features.

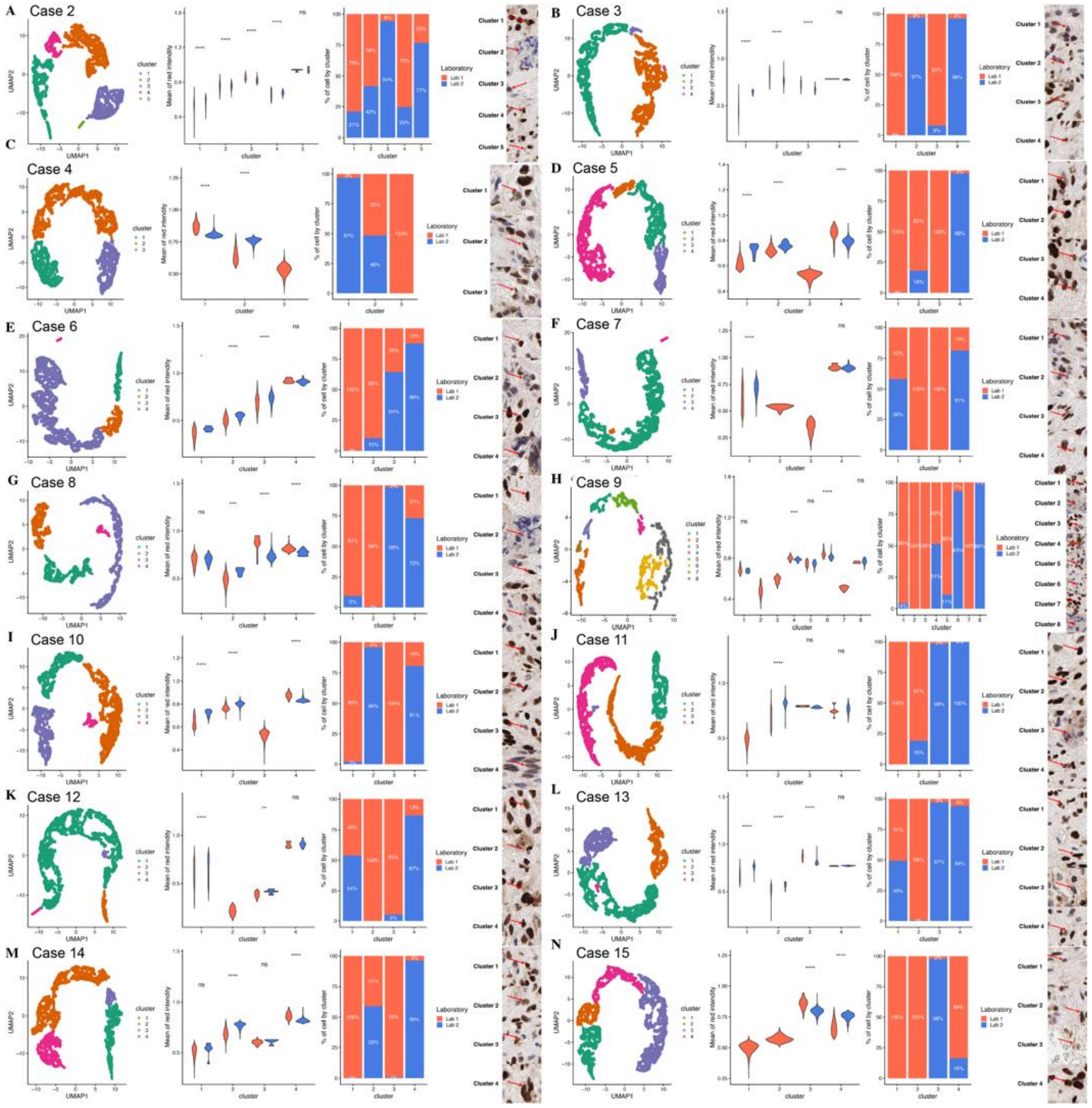


**Supplementary Figure 2: Analysis of Tissue Cell Stanning for Inter-Laboratory Variability in Image Processing and Feature Extraction.** (A-N) UMAP plot presents the DBscan cluster distribution of intensity feature, on the left showing the cluster distributions, violin plots in the middle displaying the intensity differences between clusters, barplots showing the percentage of cell distributed in each cluster and representative cell images on the right for each cluster.

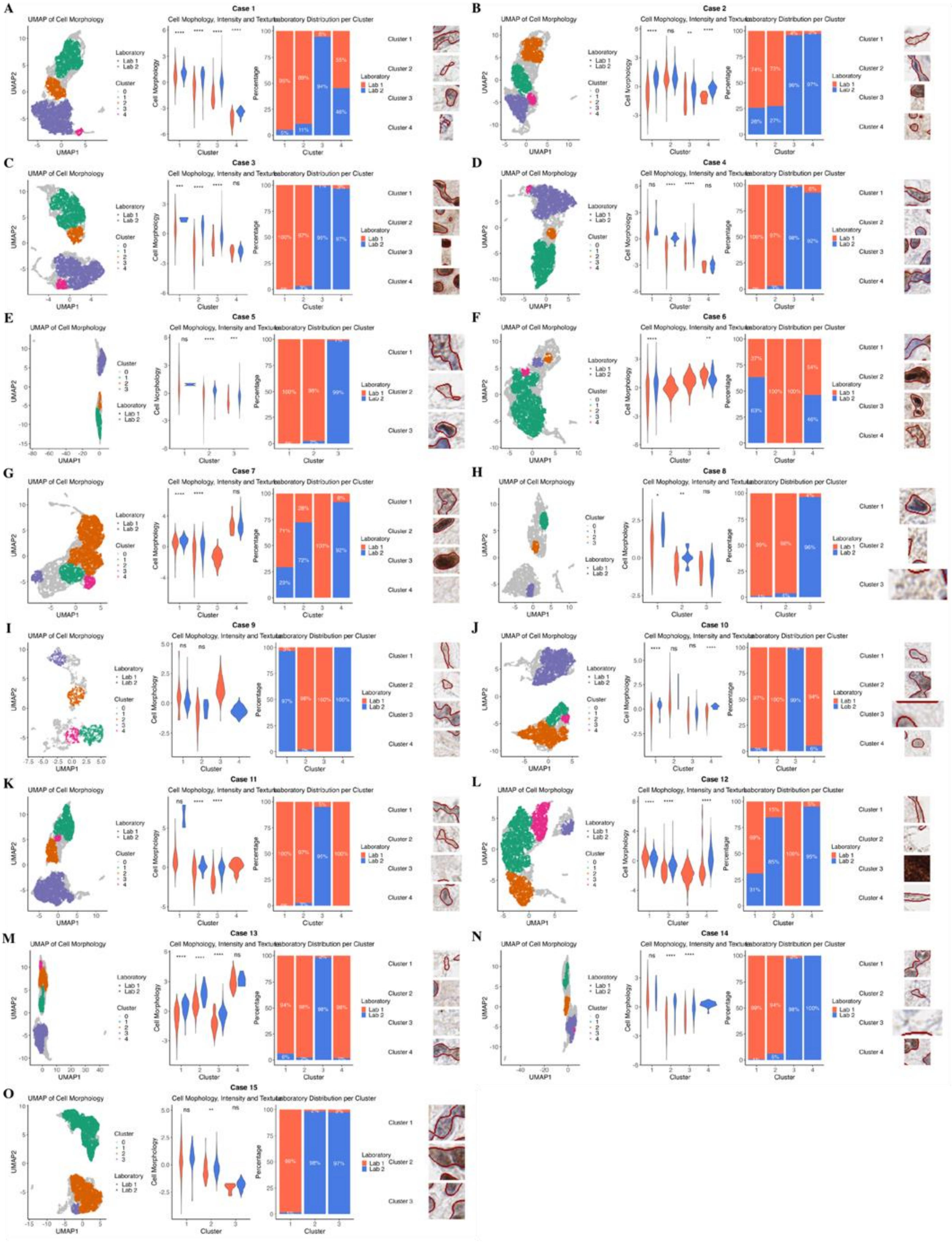
A
Case 1
UMAP of Cell Morphology
Cell Mophology, Intensity and Texture
Laboratory Distribution per Cluster
B
Case 2
C
Case 3
D
Case 4
E
Case 5
F
Case 6
G
Case 7
H
Case 8
I
Case 9
J
Case 10
K
Case 11
L
Case 12
M
Case 13
N
Case 14
O
Case 15
UMAP1
UMAP2
Cell Morphology
Percentage
Cluster
Laboratory
Lab 1
Lab 2
Cluster 1
Cluster 2
Cluster 3
Cluster 4

**Supplementary Figure 3: Analysis of Tissue Cell Stanning for Inter-Laboratory Variability in Image Processing and Feature Extraction.** (A-N) UMAP plot presents the DBscan cluster distribution of morphology: intensity, texture, shape, position and density features, on the left showing the cluster distributions, violin plots in the middle displaying the morphology differences between clusters, barplots showing the percentage of cell distributed in each cluster and representative cell images on the right for each cluster.

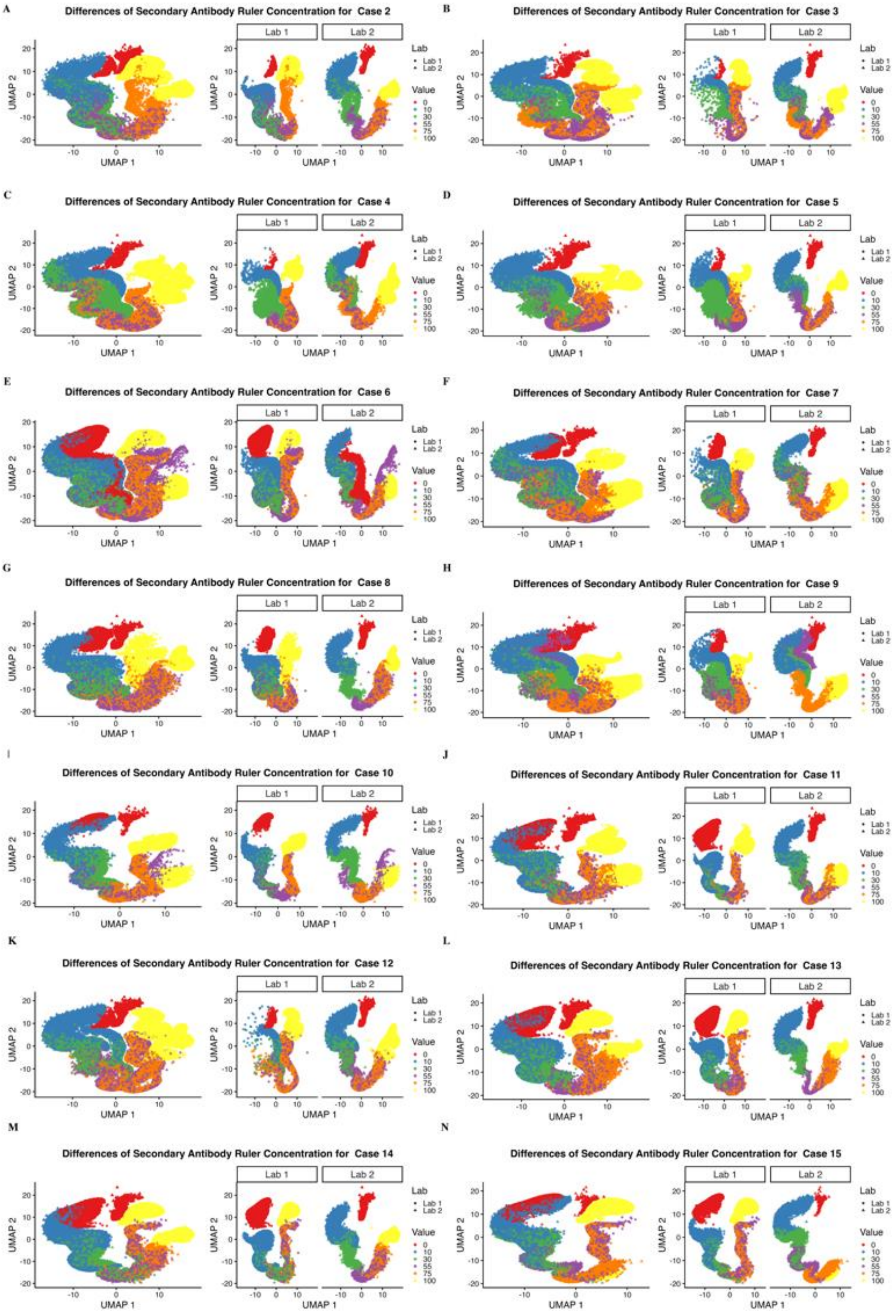
A
Differences of Secondary Antibody Ruler Concentration for Case 2
B
Differences of Secondary Antibody Ruler Concentration for Case 3
C
Differences of Secondary Antibody Ruler Concentration for Case 4
D
Differences of Secondary Antibody Ruler Concentration for Case 5
E
Differences of Secondary Antibody Ruler Concentration for Case 6
F
Differences of Secondary Antibody Ruler Concentration for Case 7
G
Differences of Secondary Antibody Ruler Concentration for Case 8
H
Differences of Secondary Antibody Ruler Concentration for Case 9
I
Differences of Secondary Antibody Ruler Concentration for Case 10
J
Differences of Secondary Antibody Ruler Concentration for Case 11
K
Differences of Secondary Antibody Ruler Concentration for Case 12
L
Differences of Secondary Antibody Ruler Concentration for Case 13
M
Differences of Secondary Antibody Ruler Concentration for Case 14
N
Differences of Secondary Antibody Ruler Concentration for Case 15
Lab 1
Lab 2
UMAP 1
UMAP 2
Lab
Value
0
10
30
55
75
100

**Supplementary Figure 4: Analysis of Secondary Antibody Concentration Ruler for Inter-Laboratory Variability in Image Processing and Feature Extraction.** (A–N) UMAPs showing differences in the distribution of each secondary antibody concentration across laboratories for each case.

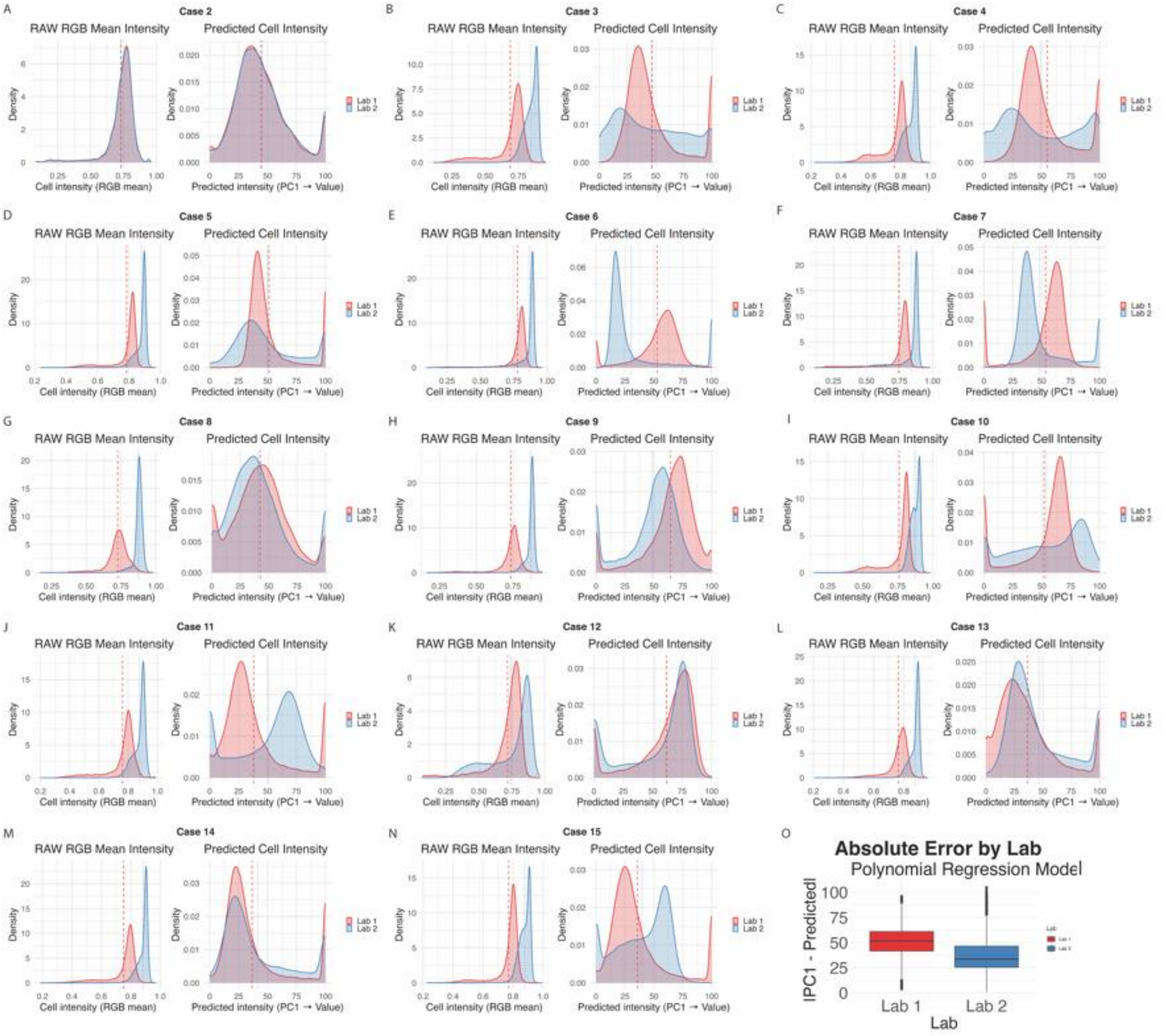


**Supplementary Figure 5. Predictive model for quantifying p53 antibody cell intensity between two laboratories (Lab 1 and Lab 2).** (A-N) Histogram from Cases demonstrating the harmonization effect. (O) The boxplots display Comparison of measurement distributions.

**Supplementary Table 1**: Percent reduction of the per-case inter-laboratory calibration technology Δ|SMD| before and after harmonization,

| ***Case*** | ***\|SMD\| RAW*** | ***\|SMD\| PRED*** | $\Delta\|SMD\|$ | ***% reduction*** |
|---|---|---|---|---|
| ***Case_1*** | 2.12883576 | 0.16119591 | 1.96763986 | 92.4279783 |
| ***Case_2*** | 0.00224944 | 0.07791183 | -0.0756624 | NA |
| ***Case_3*** | 1.7402548 | 0.1723174 | 1.5679374 | 90.0981511 |
| ***Case_4*** | 1.65998441 | 0.28504992 | 1.37493449 | 82.8281566 |
| ***Case_5*** | 1.56100372 | 0.18856122 | 1.3724425 | 87.9205144 |
| ***Case_6*** | 1.12046263 | 0.12972931 | 0.99073331 | 88.421808 |
| ***Case_7*** | 0.88537675 | 0.06775231 | 0.81762443 | 92.3476289 |
| ***Case_8*** | 2.63437984 | 0.11006772 | 2.52431212 | 95.8218735 |
| ***Case_9*** | 2.33827232 | 0.40614445 | 1.93212787 | 82.6305754 |
| ***Case_10*** | 1.64714308 | 0.12796529 | 1.51917779 | 92.231076 |
| ***Case_11*** | 1.58492265 | 0.18065309 | 1.40426956 | 88.6017726 |
| ***Case_12*** | 0.17557956 | 0.19922754 | -0.023648 | -13.468526 |
| ***Case_13*** | 2.01893629 | 0.28527256 | 1.73366373 | 85.8701556 |
| ***Case_14*** | 1.90642958 | 0.10431353 | 1.80211605 | 94.5283305 |
| ***Case_15*** | 1.77204265 | 0.07314456 | 1.69889808 | 95.8723025 |
| ***Total*** | 1.66 | 0.16 | 1.5 | 90.3 |

RAW: PC1 before prediction in pixels units; PRED: predicted after calibrated in %; |SMD|: Standard Mean Deviation; Δ|SMD| $= (RAW - PRED)$; NA: data < 0,10 threshold.
